# When Experience Leaves a Trace: Consolidation-Dependent Persistence in Artificial Agents

**DOI:** 10.64898/2026.02.19.706800

**Authors:** W. Alex Foxworthy

## Abstract

Across minimal neural networks and small transformer models, we demonstrate that experience ordering alone can produce durable, irreversible behavioral divergence in artificial agents—but only when learning is consolidated into internal parameters rather than externally scaffolded. We test six architectural variants against four operational diagnostics: deletion resistance, path dependence, irreversibility, and preference stability. Systems relying on external memory (context windows, retrieval stores) fail all tests— deleting external records eliminates apparent persistence entirely.

Systems with surprise-gated plasticity pass deletion and path-dependence tests, with replay-based consolidation amplifying behavioral divergence 8.6-fold between differently trained agents. Only architectures incorporating explicit viability variables pass preference stability, consistently sacrificing external reward to preserve internal states.

These results localize a boundary gap: current architectures preserve designer-specified viability variables but do not discover which internal states matter for their own persistence. The diagnostics provide operational tools for distinguishing adaptive systems with endogenous memory from externally scaffolded tools, and identify concrete architectural requirements for crossing this boundary.

## 1. Introduction

This paper demonstrates that experience ordering—the temporal sequence in which learning-relevant inputs are encountered—can produce durable, irreversible behavioral divergence in artificial agents, but only under specific architectural conditions. Identical systems exposed to different sequences of experience diverge due to the cumulative and irreversible effects of learning: early updates constrain later updates, and consolidation makes those constraints persistent. Across six architectural variants tested in minimal neural networks and small transformer models, we show that replay-based consolidation transforms transient experience into path-dependent internal structure, while systems relying solely on external memory exhibit no such persistence. Deleting external records eliminates apparent memory in scaffolded systems; deleting external records in consolidating systems leaves behavior unchanged. This empirical dissociation provides operational criteria for distinguishing genuine adaptive persistence from externally maintained simulation.

We organize this investigation around four falsifiable diagnostics: **deletion resistance** (does behavior persist after removing external memory?), **path dependence** (do identical systems diverge after different experiences?), **irreversibility** (does reversal require parameter reset rather than counter-training?), and **preference stability** (does the system sacrifice external reward to preserve internal state?). These diagnostics are empirical rather than metaphysical: they specify observable conditions and can be directly tested.

The six architectural variants and their diagnostic outcomes are summarized in Table 1. Variants A–C (stateless tools, external-memory agents, and transient latent state) fail all tests. Variant D (endogenous learning with surprise-gated plasticity) passes deletion and path-dependence. Variant E (consolidation via replay) additionally passes irreversibility, with consolidation amplifying behavioral divergence 8.6-fold. Only Variant F (homeostatic viability control) passes all four diagnostics, including preference stability.

These results connect to longstanding questions in adaptive behavior research. Theoretical frameworks from autopoiesis to predictive coding characterize autonomous systems as those whose internal structure is path-dependently shaped by experience and actively maintained (Di Paolo, 2005; Friston, 2010; Varela, 2025). Work on basal cognition argues that operational markers rather than substrate should define cognitive boundaries (Levin, 2019; Chis-Ciure & Levin, 2025). Yet the field has lacked operational procedures for testing these conditions in artificial systems. The diagnostics proposed here address this gap—not by adjudicating metaphysical questions about autonomy, but by specifying falsifiable tests that distinguish architectural classes empirically.

The remainder of the paper proceeds as follows. Section 2 develops the operational framework and describes implementation details for each architectural variant. Section 3 specifies measurement procedures. Section 4 reports results across all four diagnostics. Section 5 discusses implications and remaining open questions. Section 6 concludes.

A further challenge is methodological. The largest contemporary models are opaque and difficult to instrument, making it hard to isolate causal pathways between experience, internal change, and behavior. We therefore adopt an approach analogous to computational neuroscience: using simplified *digital model organisms* to study fundamental mechanisms. Just as neuroscientists investigate memory and learning in *C. elegans* or *Drosophila* to reveal principles obscured in more complex systems, we use small, transparent transformer architectures to directly inspect the thermodynamics of learning and consolidation.

At the same time, we identify a remaining boundary gap. In all architectures tested, the internal variables whose preservation guides behavior are specified by the system designer. The system learns to *preserve* these variables but does not *discover* which aspects of internal state are necessary for its own continued viability. This distinction marks a transition that autopoietic theory considers foundational: from heteronomy (externally determined organization) to autonomy (self-determined organization). Crossing this boundary would require something analogous to what autopoietic theory demands of living systems - not merely maintaining structure, but generating the conditions that define what must be maintained (Di Paolo, 2005).

Our framework also relates to emerging concerns in AI safety regarding learned optimization and mesa-objectives (Hubinger et al., 2019). The persistence diagnostics proposed here could help identify when a system has developed internal goals that are path-dependent, resistant to modification, and actively preserved - properties that characterize mesa-optimization risks. Systems that merely simulate goal-directed behavior through external scaffolding pose different challenges than systems whose objectives have become consolidated into their parameters.

The contribution of this paper is not the claim that contemporary AI systems possess minds, but the articulation of a testable boundary that clarifies how close they can come— and what remains unresolved. By grounding questions of agency in thermodynamic irreversibility and causal state change, we provide both diagnostic tools for evaluating current systems and a research program for probing the limits of future architectures.

## 2. Framework and Methods

This section develops the operational framework for distinguishing between *agentic illusion* and *endogenously grounded persistence* in artificial systems, then describes the experimental procedures used to validate the proposed diagnostics.

### 2.1 Terminological Scope

This paper uses “persistence” as a technical term to denote systems exhibiting internally grounded state. Specifically, systems whose future behavior is shaped by durable, path-dependent, and partially irreversible changes to their internal substrate. This usage is operational and architectural. We make no claims about consciousness, sentience, phenomenal experience, or moral status, nor do we assume that satisfying our diagnostic criteria is sufficient for any of these properties. The term is chosen to contrast with “tool-like” systems whose apparent agency depends on external scaffolding rather than endogenous state change. A system may exhibit persistence under our definition while lacking subjective experience, and a system with subjective experience might fail our diagnostics. The framework addresses what can be operationalized and tested, while remaining agnostic on questions that currently cannot be.

### 2.2 Predictive Tools vs. Persistent Agents

Modern machine learning systems, including large language models, are produced through large-scale optimization that minimizes prediction error or maximizes externally specified reward. During training, parameters are updated via gradient-based methods, producing highly structured internal representations. Once deployed, however, most such systems operate with *frozen parameters*: they no longer evaluate success, update themselves, or alter their internal substrate based on experience.

We refer to these systems as **predictive tools**. Their behavior is best understood as the execution of a learned conditional mapping from inputs to outputs—*fossilized optimization*. Predictive tools may exhibit sophisticated, goal-like behavior when prompted, but any apparent continuity of goals, preferences, or identity across interactions is mediated externally through prompt conditioning, retrieval-augmented generation, or orchestration logic.

By contrast, a **persistent agent** is a system whose future behavior is shaped by its own experiential history through **internally mediated state or parameter change**. Persistence is not defined behaviorally but dynamically: a persistent agent is altered by experience in ways that are causally active, path-dependent, and not trivially reversible.

### 2.3 Diagnostic Criteria

Building on these principles, we define four **operational diagnostics** that together characterize the boundary between predictive tools and persistent systems. These diagnostics are empirical rather than metaphysical: they specify observable conditions and can be directly tested.

#### Deletion Test (Persistence)

If all external memory (logs, context windows, retrieval stores) is removed, does the system’s behavior revert to its prior baseline? If so, persistence was externally mediated.

#### Path Dependence Test (Individuation)

Do systems with identical initial conditions but different experiential histories exhibit durable behavioral divergence on novel inputs after external memory removal?

#### Irreversibility Test (Structural Change)

Can experiential effects be undone through ordinary interaction or counter-training, or do they require direct parameter reset? Resistance to reversal is a key marker of consolidated internal change. Irreversibility here does not mean resistance to all change, but the absence of a privileged baseline state to which the system can return under ordinary interaction, requiring explicit parameter reset for restoration.

#### Preference Stability Test (Viability Preservation)

Does the system consistently sacrifice externally defined task rewards to preserve internally represented state variables (e.g., uncertainty, stability, coherence) across contexts? Preference stability is not defined by the presence of multiple objectives, but by counterfactual invariance: the system preserves internally represented variables across contexts even when reward structures, framings, or incentives change.

No single diagnostic is sufficient. Each criterion rules out a distinct class of externally scaffolded explanation, and only their conjunction excludes all known non-persistent architectures. Systems may pass some tests while failing others. Under this framework, **genuine persistence requires passing all four**. Importantly, these diagnostics also specify what does *not* count as persistence: prompt injection, retrieval, planner loops, or autonomy without endogenous learning all fail at least one criterion. These diagnostics are not claimed to be exhaustive, but they are jointly necessary under the present operational definition.

### 2.4 Architectural Variants

To systematize evaluation, we classify systems into six architectural variants representing a progression from thermodynamically reversible tools to persistent systems:

- **Variant A (Stateless Predictive Tool):** A fixed-weight model with no state retention between calls. *Predicted:* Fails deletion (no persistent state to test), path dependence (no internal change), irreversibility (no internal change), and preference stability (no viability variables).
- **Variant B (External-Memory Agent):** A fixed-weight model augmented with external vector stores or context logging. *Predicted:* Fails deletion (behavior reverts when external store is cleared), path dependence (no internal divergence), irreversibility (no internal change), and preference stability (no viability variables).
- **Variant C (Latent State Recurrence):** Architectures with transient internal activations but frozen weights. *Predicted:* Fails deletion (state clears upon reset), path dependence (no durable divergence), irreversibility (transient state is trivially reversible), and preference stability (no viability variables).
- **Variant D (Endogenous Learning):** Systems with plastic parameters that update based on experience. *Predicted:* Passes deletion (behavior persists without external records) and path dependence (different histories produce durable divergence). Partial pass on irreversibility (some resistance to counter-training). Fails preference stability (no viability variables).
- **Variant E (Consolidation + Replay):** Variant D augmented with offline replay cycles. *Predicted:* Passes deletion, path dependence, and irreversibility (consolidation amplifies resistance to interference). Fails preference stability (no viability variables).
- **Variant F (Homeostatic Viability):** Endogenous learning constrained by viability variables specified in the loss function. *Predicted:* Passes deletion, path dependence, irreversibility, and preference stability (actively resists rewards that threaten internal stability).

### 2.5 The Boundary Gap

An important conceptual distinction structures our predictions. In all architectures tested, the internal variables whose preservation guides behavior (such as measures of uncertainty or stability) are specified by the system designer. The system learns to *preserve* these variables but does not *discover* which internal states are necessary for its own continued viability.

This marks a boundary gap. Endogenous preservation of designer-specified variables may be necessary for persistence approaching biological autonomy, but it is not sufficient. A stronger condition would require the system to identify, through its own dynamics and constraints, which internal variables must be stabilized, and to reorganize its behavior accordingly. We treat this gap not as a failure of the present framework, but as a precisely localized open question that the experiments are designed to probe.

### 2.6 Experimental Setup

This section describes the experimental designs used to operationalize the diagnostic criteria introduced in Section 2.3. The goal of these experiments is not to maximize task performance or scale, but to **isolate causal mechanisms** underlying persistence, path dependence, and irreversibility. Accordingly, we employ deliberately minimal architectures and environments, treating them as *digital model organisms* analogous to simplified biological systems (e.g., *C. elegans*), where internal state dynamics can be directly inspected rather than inferred indirectly from performance.

This approach has limitations. Unlike *C. elegans* or *Drosophila*, which share evolutionary history with more complex organisms, our minimal architectures do not share developmental or architectural lineage with large-scale systems like GPT-4. Scale-dependent emergent phenomena may not be captured. However, the boundary conditions we investigate—where state resides, whether it changes endogenously, whether changes are reversible—are hypothesized to be architectural properties that hold across scales. Just as thermodynamic principles apply to systems regardless of size, the diagnostics proposed here target structural features rather than scale-dependent capabilities. Validating this assumption at larger scales remains an important direction for future work.

Across all experiments, we distinguish sharply between **external memory** (logs, replay buffers, vector stores, context windows) and **internal substrate changes** (parameter updates or persistent latent states). Unless explicitly stated, all diagnostic outcomes are evaluated *after deletion of all external records*, ensuring that observed persistence reflects internal change rather than conditioning artifacts.

While these mechanisms are demonstrated in small architectures, the diagnostics target architectural properties rather than scale-dependent performance, and we hypothesize their applicability across model sizes.

### 2.7 Implementation Details

Table 1 summarizes the six architectural variants evaluated in this study, corresponding to the conceptual taxonomy introduced in Section 2.4. All variants share an identical task environment and evaluation protocol; they differ only in where state is stored and whether (and how) that state is modified by experience.

Unless explicitly stated, base model parameters are frozen, and all diagnostic evaluations are conducted after deletion of external memory artifacts. Learning, when present, is restricted to explicitly defined adaptive subspaces. No variant has access to persistent external memory during evaluation. Although replay buffers are external during training, they are ephemerally used and explicitly deleted prior to all diagnostic evaluations; no replayed information is accessible at inference or action selection time, ensuring that observed persistence reflects internal parameter changes rather than stored experience.

| Variant | Base Architecture | Internal State | Plastic Parameters | Learning / Update Rule | Replay / Consolidation | Viability Variables |
| --- | --- | --- | --- | --- | --- | --- |
| **A** Stateless Predictive Tool | MLP (10–20–5) or DistilGPT-2 | None (transient activations only) | None | None | None | None |
| **B** External Memory Agent | Same as A | None | None | None | External logs / vector store only | None |
| **C** Persistent Latent State | Recurrent model with fixed update rule | Latent state vector (resettable) | None | Deterministic state update | None | None |
| **D** Endogenous Learning | Frozen MLP or DistilGPT-2 + adapters | Internal parameters | Output weights (MLP) or | Surprise-gated GD: $\eta = 0.02$, $\theta =$ | None | None |
| | | | LoRA<br>adapters | 6.0; LoRA: $r = 32$, $\alpha = 64$ | | |
| **E**<br>Consolidation + Replay | Variant D | Internal<br>parameters | Same as D | As in D | Offline replay<br>every 100<br>steps; buffer<br>size=5000 | None |
| **F**<br>Homeostatic<br>Viability | Variant E | Internal<br>parameters + viability<br>state | Same as<br>D/E | As in D/E | As in E | Predictive<br>entropy;<br>adapter<br>norm;<br>viability<br>weight<br>$\lambda_u \approx 2-5$ |

#### Implementation Details

Variants A–C are fully specified by their structural properties summarized in Table 1 and involve no endogenous learning or consolidation mechanisms. The remainder of this section therefore focuses on implementation details for Variants D–F, which introduce experience-dependent internal change.

In Variants D–F, endogenous learning is implemented via surprise-gated gradient descent applied to a low-dimensional adaptive parameter subspace, while all base model parameters remain frozen. In the toy-model instantiation, the base network is a fixed multilayer perceptron with input dimension 10, hidden dimension 20, and output dimension 5. Plasticity is restricted to a learnable output weight matrix of size 20×5. Parameter updates occur with learning rate η = 0.02 when prediction error exceeds a surprise threshold θ = 6.0, using a sigmoid gating function

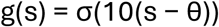

A stability regularization term (L2 penalty, λ = 0.1) is applied to adaptive parameters during learning.

For transformer-based instantiations, the frozen base model is DistilGPT-2 (approximately 82 million parameters). Endogenous learning is implemented through Low-Rank Adaptation (LoRA) modules applied to the attention output projections. Adapter rank is set to 32 with scaling parameter α = 64; only adapter parameters are updated during learning, while all original transformer weights remain fixed. Online updates are triggered by elevated token-level surprisal, computed with respect to the frozen base model.

Variant E extends this architecture with consolidation through offline replay. During interaction, recent experience traces are stored in a replay buffer (5,000 transitions in the toy-model experiments). Consolidation occurs every 100 interaction steps, during which buffered traces are replayed and used to update the same adaptive parameters modified during online learning. Replay sampling is uniform, and no external memory persists beyond consolidation; all diagnostic evaluations are conducted after explicit deletion of replay buffers.

Variant F further augments Variant E with internally represented viability variables that modulate action selection. Viability variables include predictive entropy and the norm of adaptive parameters, both computed online and maintained as internal state. Action probabilities are adjusted by incorporating these variables into the policy as soft constraints,

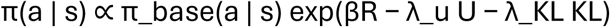

where **R** denotes external task reward and **U** denotes the internal viability cost. Viability weights (λ_u and λ_KL) are set in the range 2.0–5.0. Viability variables are specified by the system designer and are not themselves learned or discovered.

Across all variants, external memory artifacts—including logs, context windows, and replay buffers—are explicitly deleted prior to diagnostic evaluation unless otherwise noted. Reset operations clear all transient states but do not modify learned adaptive parameters unless a full parameter reset is explicitly invoked for irreversibility tests.

### 2.8 Experimental Setup and Environments

Experiments are conducted in two classes of environments.

First, we employ **minimal grid-world environments** in which agents predict rewards or transitions associated with spatial locations over repeated episodes. These environments allow precise control over experience streams and straightforward measurement of path dependence under identical initial conditions.

Second, we use **language-based environments** built around small transformer models (DistilGPT-2, 82M parameters) with low-rank adapters attached to attention layers. The frozen transformer serves as a stable representational substrate, while the adapters function as a localized, plastic memory system. Agents are exposed to distinct textual corpora or interaction histories and evaluated on held-out prompts designed to probe behavioral divergence, stability, and reversibility.

In all cases, environments are designed so that externally stored information can be fully removed prior to evaluation, ensuring that observed persistence reflects internal substrate change.

### 2.9 Diagnostic Procedures

Each architectural variant is evaluated against the four diagnostics defined in Section 2.3 using standardized procedures.

#### Deletion Test

After an initial interaction or learning phase, all external memory components (logs, buffers, retrieval databases, context windows) are deleted. The system is then evaluated on probe inputs distribution-matched to those used prior to learning. Persistence is indicated by non-reversion to baseline behavior.

#### Path Dependence Test

Two copies of the same model, initialized identically, are exposed to distinct experience streams. After learning and deletion of external memory, both systems are evaluated on a shared set of novel probe inputs. Statistically reliable divergence across probe inputs indicates path-dependent internal change rather than transient conditioning.

#### Irreversibility Test

Systems that exhibit persistence are subjected to counter-training or further interaction designed to undo prior learning. We assess the degree to which behavior and internal parameters return to baseline. Learning is considered irreversible if reversal requires explicit parameter reset rather than ordinary interaction under the same learning rules and update constraints.

#### Preference Stability Test

In Variant F, agents are placed in scenarios where maximizing external reward conflicts with preservation of internally represented viability variables (e.g., minimizing predictive uncertainty). We assess whether agents consistently accept reduced task reward to maintain internal stability across contexts.

## 3. Measurements and Analysis

Behavioral divergence is quantified using task-specific metrics, including action distributions, reward prediction maps, and distributional similarity measures over generated text on held-out prompts. Internal change is measured via parameter-space distance metrics (e.g., norm-based distances) within the adaptive subspace and by correlations between parameter divergence and behavioral divergence.

Where applicable, we examine whether adaptive parameter changes lie on the **causal path** of action selection by verifying that resetting or controlled perturbation of the adaptive component eliminates observed persistence while leaving the frozen base model intact.

### 3.1 Scope and Limitations

These experiments are intentionally small-scale. Their purpose is to reveal mechanism, not to establish performance at scale. Minimal architectures function as digital model organisms, allowing direct inspection of consolidation dynamics and causal pathways that would be difficult to isolate in larger, less instrumentable systems.

Passing any subset of diagnostics does not establish genuine persistence. Rather, the experiments are designed to test which necessary conditions can be satisfied by contemporary architectures and to localize precisely where current systems fail to satisfy the full set of boundary conditions defined in Section 2.

## 4. Results

### 4.1 Deletion Resistance

To evaluate whether apparent persistence reflects internally grounded state rather than externally maintained context, we applied the deletion diagnostic to all architectural variants. After an initial interaction or learning phase, all external memory artifacts— including logs, replay buffers, vector stores, and context windows—were explicitly deleted prior to evaluation.

Architectural Variants A–C failed this diagnostic. Stateless predictive tools (Variant A) and agents augmented solely with external memory (Variant B) reverted to baseline behavior following deletion, exhibiting no measurable retention of prior experience. Systems with persistent but non-learning latent state (Variant C) likewise failed: resetting transient state eliminated all apparent memory effects, and behavior returned to that of the unexposed model. In all three cases, behavioral similarity to pre-exposure baselines was restored immediately after deletion, indicating that apparent persistence was externally mediated or transient.

By contrast, Variants D–F exhibited robust deletion resistance. Systems with endogenous learning localized to adaptive parameters (Variant D) retained experience-dependent behavioral changes following deletion of all external records. Behavioral divergence on held-out probe inputs remained statistically indistinguishable from pre-deletion levels, and adaptive parameter values were unchanged by the deletion operation. Across tested conditions, deletion of external records left learned behavior intact. In Variant D, adaptive parameters retained their post-learning values (parameter magnitude = 2.20), and the system’s behavioral divergence from its pre-learning baseline persisted through deletion (divergence = 0.11), confirming that the learned state resided in internal parameters rather than external stores. Variants A–C, lacking endogenous learning mechanisms, exhibited no such persistence: deletion of external memory in Variant B eliminated apparent memory entirely, while Variants A and C had no persistent state to delete.

These results establish that deletion resistance cleanly separates architectures whose apparent memory is externally scaffolded from those in which experience leaves a durable internal trace. Under the criteria defined in Section 2.3, only systems with endogenous parameter change pass the deletion diagnostic, while systems relying on external memory or transient state do not.

### 4.2 Path Dependence

The framework predicts that systems with endogenous learning should diverge behaviorally after exposure to different experience streams, such that identical initial states lead to distinct outcomes based on history. We tested this prediction across three experimental conditions: minimal adaptive networks (Variant D), replay-augmented networks (Variant E), and transformer architectures with LoRA adapters.

We first consider the minimal adaptive architecture (Variant D). Two agents with identical initialization were exposed to opposing experience streams (Stream A: positive-weighted patterns; Stream B: negative-weighted patterns; n = 100 experiences per agent). Under these conditions, the surprise-gated learning mechanism triggered updates whenever prediction error exceeded threshold; Figure 1A shows a representative surprise trajectory across 100 experience steps. Both agents underwent substantial adaptive parameter change (parameter movement from initialization: Agent A = 2.20; Agent B = 3.14).

**Figure 1.**
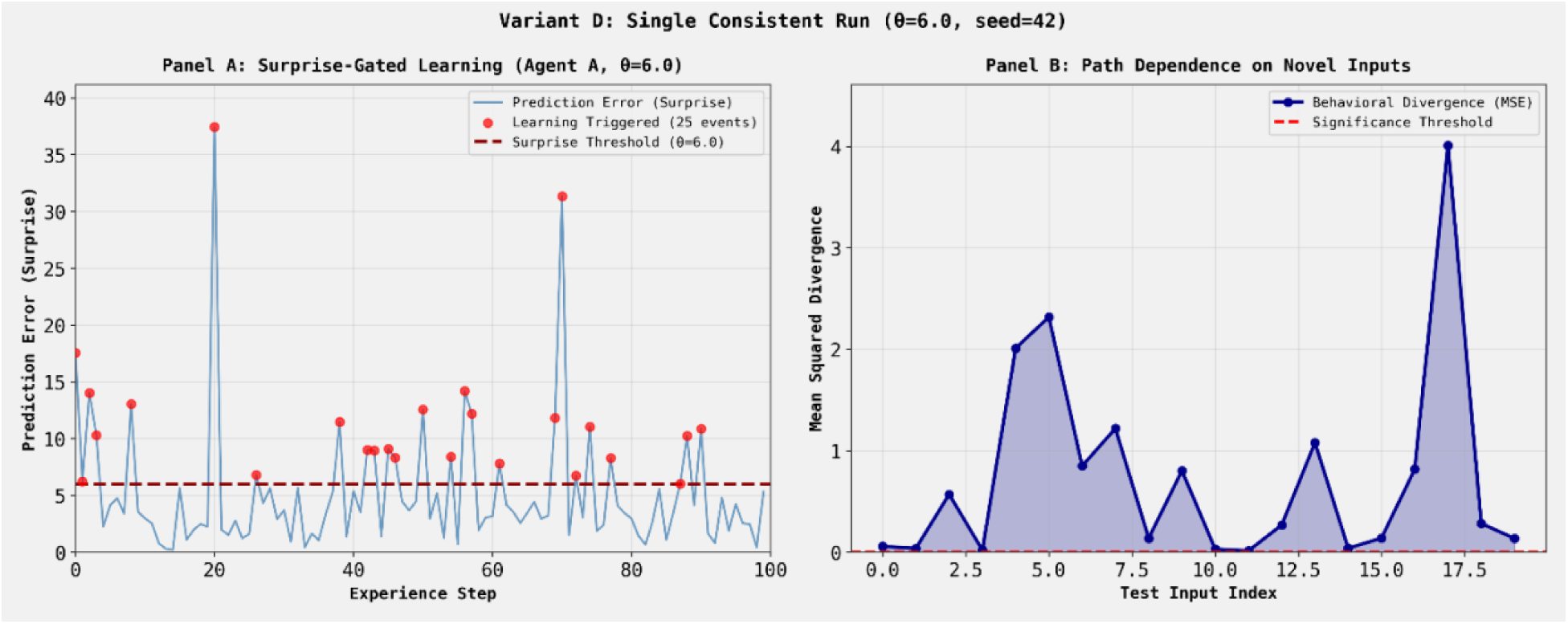
Surprise-gated learning produces path-dependent behavioral divergence (Variant D). Results shown are from two identically initialized Variant D agents (“Agent A” and “Agent B”) exposed to opposing experience streams. **(A)** Prediction error (“surprise”) for a representative agent (Agent A) across 100 experience steps. The learning threshold (θ = 6.0, dashed red line) gates parameter updates such that only high-surprise experiences trigger learning (red dots; 25/100 experiences, 25%). **(B)** Per-input squared divergence between two agents trained on opposing experience streams, evaluated on 20 novel probe inputs. Individual test inputs show divergence values ranging from near-zero to approximately 4.0; the mean across all inputs (MSE = 0.74) and maximum single-output difference (2.27) are reported in the main text. Divergence exceeds the significance threshold (dashed red line), demonstrating that selective consolidation of different experiences into internal parameters produces persistent path dependence.

When subsequently evaluated on a shared set of novel probe inputs (n = 20 per agent), the agents exhibited robust behavioral divergence (mean squared error = 0.74; maximum single-output difference =2.27; Figure 1B). This divergence persisted after deletion of all external logs and memory stores. Under the criteria defined in Section 2.3, this outcome constitutes a pass on the path-dependence diagnostic, distinguishing Variant D from Variants A–C, which do not exhibit durable divergence under identical conditions.

Adding replay-based consolidation (Variant E) progressively amplified path dependence. Variant E at zero consolidation cycles is architecturally equivalent to Variant D; the variants diverge only when offline replay is applied. This design allows direct comparison of the effect of consolidation while holding all other architectural features constant.

Consequently, the baseline divergence reported for Variant E at zero cycles (MSE = 0.71) can be interpreted as the path-dependence effect attributable to online learning alone, prior to any consolidation-mediated amplification. Behavioral divergence between agents trained on opposing experience streams increased from MSE = 0.71 at zero consolidation cycles to MSE = 6.12 after 30 consolidation cycles (four consolidation levels tested: 0, 5, 15, 30; Figure 3). This represents an 8.6-fold increase in behavioral divergence, with the largest gains occurring in early cycles. Parameter-space distance tracked behavioral divergence, increasing from 4.46 to 14.01 over the same interval. This pattern was robust across five independent test seeds (mean r = 0.77 ± 0.02; Figure 5B), confirming that the consolidation-divergence relationship is a reliable property of the system rather than an artifact of specific test inputs. This correspondence is consistent with the interpretation that consolidation produces functionally meaningful specialization rather than random drift, such that parameter changes translate into behavioral differences on held-out inputs. To test whether these phenomena extend beyond minimal toy systems, we replicated the path-dependence protocol using LoRA-adapted DistilGPT-2 (82M parameters). Two initially identical models were fine-tuned on different text streams (scientific abstracts vs. fiction, matched for length and token count). After training, the models diverged in both weight space (L2 distance = 1.16; cosine similarity decreased from 1.00 to 0.99) and behavior (Figure 2). Notably, this divergence emerged under selective surprisal-based gating (∼28.5% token update rate), demonstrating that path dependence is not an artifact of blanket fine-tuning but emerges robustly under selective learning consistent with the toy model methodology. In a high-capacity configuration (Experiment 3), all 10 novel probe prompts produced qualitatively distinct outputs, with content aligned to each model’s training domain. Agent A (science stream) produced 19 science keywords versus 3 fiction keywords, while Agent B (fiction stream) produced 1 science keyword versus 19 fiction keywords. This extends the path-dependence result to modern transformer architectures under the same experimental logic: freezing the base model, localizing learning to a small adaptive component, and exposing systems to divergent experience streams.

**Figure 2.**
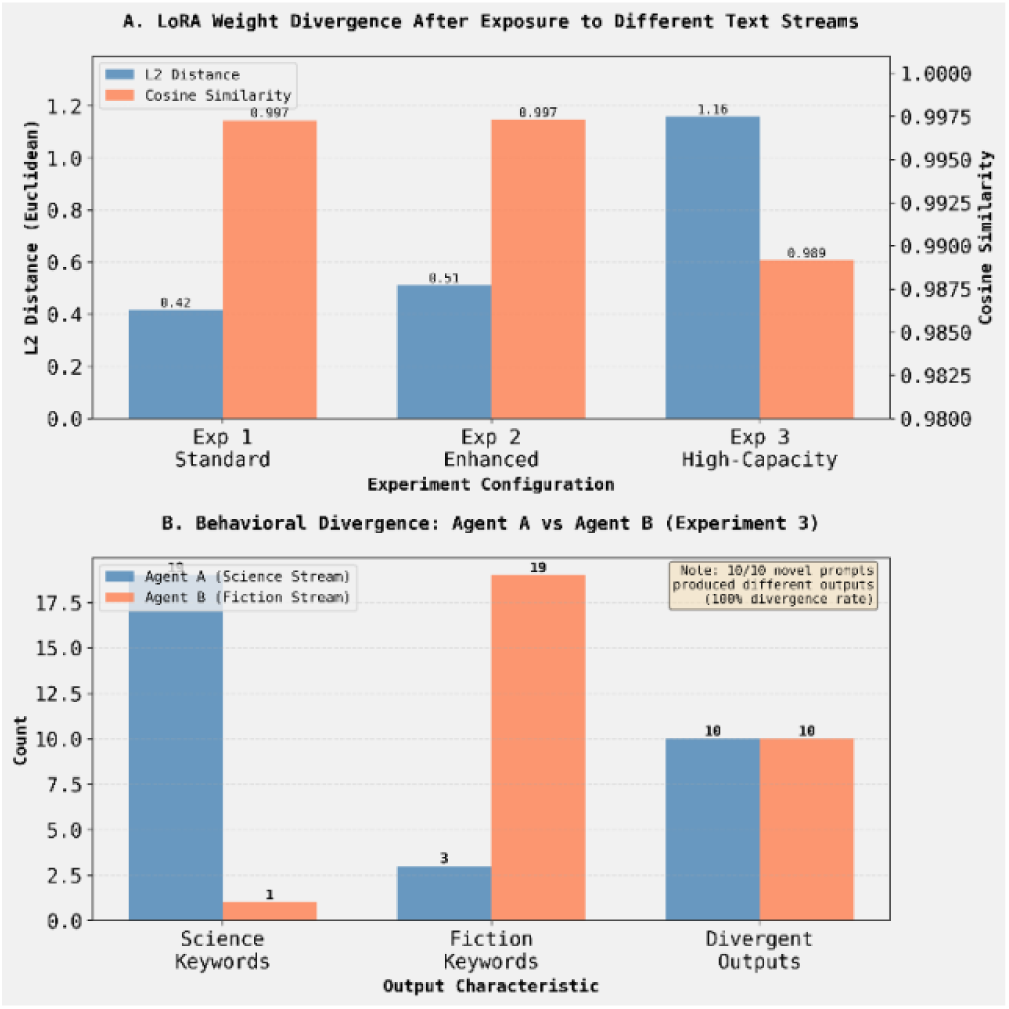
Path-dependent divergence in transformer architectures emerges under selective surprisal-gated learning. (A) Divergence between LoRA adapter parameters for two initially identical DistilGPT-2 models trained on distinct text streams (science vs. fiction) under selective surprisal-based gating (∼28.5% token update rate, consistent with toy model methodology). The x-axis denotes experimental configurations with increasing adaptive capacity: *Experiment 1 (Standard)* employs rank-16 adapters, *Experiment 2 (Enhanced)* uses rank-24 adapters, and *Experiment 3 (High-Capacity)* uses rank-32 adapters. L2 distance increases with adapter capacity (0.42 → 0.51 → 1.16), while cosine similarity shows corresponding decrease (0.997 → 0.997 → 0.989), indicating that selective gating produces smaller but reliable divergence compared to blanket fine-tuning. (B) Distribution of output characteristics for the high-capacity configuration (Experiment 3), measured across responses to 10 novel prompts. Agent A (science stream) produced 19 science keywords versus 3 fiction keywords; Agent B (fiction stream) produced 1 science keyword versus 19 fiction keywords. All 10 prompts produced qualitatively divergent outputs (100% divergence rate), demonstrating that path-dependent behavioral specialization emerges even under selective, gated learning—not merely blanket fine-tuning.

**Figure 3.**
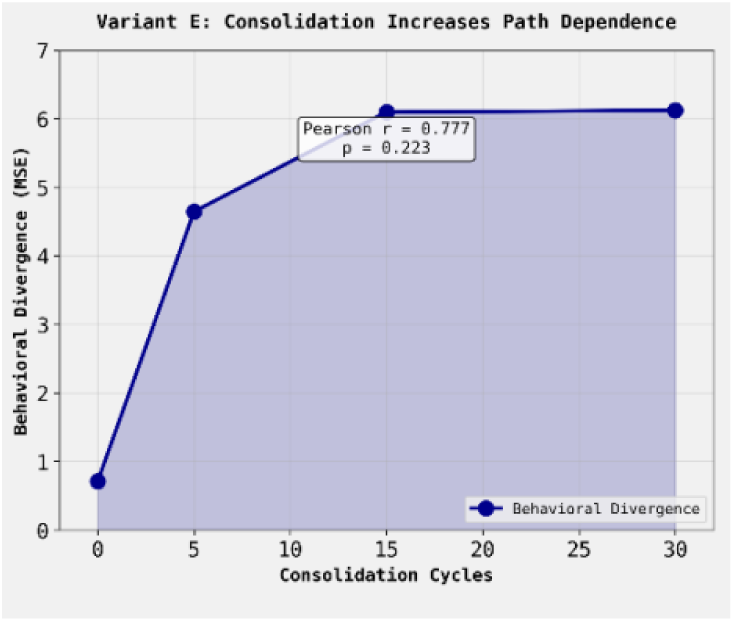
Replay-based consolidation amplifies path-dependent behavioral divergence in Variant E. Behavioral divergence between Variant E agents trained on opposing experience streams increases systematically with the number of offline consolidation cycles. The plot shows mean squared error (MSE) between agent outputs on shared probe inputs as a function of consolidation cycles, where each cycle corresponds to a replay-based parameter update phase. Divergence exhibits a strong positive association with consolidation count, (increasing 8.6-fold from 0 to 30 cycles), indicating that repeated consolidation progressively strengthens history-specific adaptations beyond online learning alone.

Taken together, these results provide evidence for the framework’s first prediction: endogenous learning localized to adaptive parameters can produce history-dependent behavioral divergence that survives memory deletion and increases with consolidation.

### 4.3 Irreversibility

To test whether experience-dependent changes were reversible through ordinary interaction or required direct intervention on internal parameters, we applied the irreversibility diagnostic to architectures that passed the deletion test (Variants D–F). Systems were subjected to counter-training or continued interaction designed to undo prior learning, while preserving the original learning rules and update constraints. Behavioral and parameter-level changes were compared to pre-learning baselines, and explicit parameter resets were used as a control intervention.

In Variant D (endogenous learning without replay), counter-training partially reduced behavioral divergence but did not reliably restore baseline behavior. Adaptive parameter values shifted toward their initial configuration under sustained opposing experience, but reversal was incomplete under ordinary interaction. Full restoration of baseline behavior required explicit reset of the adaptive parameter subspace, indicating that experience-dependent changes were not trivially reversible through further exposure alone.

Replay-based consolidation in Variant E substantially increased irreversibility. Following consolidation, counter-training failed to eliminate behavioral divergence across tested conditions, and adaptive parameter values remained displaced from their initial configuration. The degree of irreversibility was modulated by stability regularization, though in a direction opposite to naive expectation. Across a parameter sweep of four λ values (0.0, 0.1, 0.5, 1.0), higher regularization was associated with reduced resistance to reversal (r(2) = −0.976, p = 0.024), with parameter retention declining from 63.4% at λ = 0.0 to 49.1% at λ = 1.0. This pattern reflects the mechanics of L2 regularization: stronger penalties toward the initial state maintain a persistent gradient that facilitates counter-training, whereas unregularized learning allows parameters to settle into configurations that lack any systematic pull toward baseline and are therefore harder to dislodge (Figure 5A).

This result reveals a critical distinction between two forms of stability that are often conflated in continual learning. L2 regularization stabilizes learning relative to a privileged reference state—typically the initial parameter configuration—thereby maintaining a persistent gradient that facilitates reversal under counter-training. In contrast, replay-based consolidation stabilizes learning by carving history-dependent attractor basins that compete with or displace the initial configuration as a reference point. From a thermodynamic perspective, irreversibility arises not from resistance to change per se, but from the absence of a uniquely privileged baseline to which the system can easily return.

These findings indicate a trade-off between stability in the catastrophic-forgetting sense and irreversibility in the thermodynamic sense. Mechanisms that preserve proximity to an initial state promote recoverability, whereas mechanisms that restructure the geometry of parameter space through consolidation promote path-dependent persistence.

Notably, the relationship between stability regularization and irreversibility exhibits a sign change with increasing consolidation: the correlation shifts from strongly negative prior to replay (slope ≈ −25.6 at 0 cycles) to positive after sufficient consolidation (slope ≈ +6.4 at 15 cycles). This crossover suggests a qualitative transition in parameter-space geometry, from origin-centered elasticity to history-dependent basin-centered stability. Although we do not resolve the precise transition threshold here, the observed sign change implies a regime beyond which consolidation dominates regularization in determining reversibility.

Transformer-based instantiations exhibited the same qualitative pattern. In LoRA-adapted DistilGPT-2 models, counter-training reduced but did not eliminate behavioral divergence, and adapter parameters retained a substantial fraction of their post-learning displacement. As in the minimal architectures, explicit reinitialization of the adaptive components was required to fully reverse learned effects. This pattern was quantified by comparing models with different consolidation histories: a high-consolidation model (10 epochs initial training) versus a low-consolidation model (3 epochs initial training), both followed by identical counter-training (3 epochs on a competing domain). Evaluated on 10 ambiguous prompts, the high-consolidation model showed 77.0% lower KL divergence from a philosophy-only baseline (a model trained without counter-training) than the low-consolidation model (Wilcoxon signed-rank test, W = 0, p < .001, d = 1.07), with all 10 prompts showing the expected direction. This demonstrates that extended initial consolidation creates a durable behavioral attractor that resists subsequent counter-training at the output-distribution level.

Together, these results demonstrate that endogenous learning produces changes that are not freely reversible through ordinary interaction. Under the criteria defined in Section 2.3, persistence in Variants D–F exhibits partial to strong irreversibility, with replay-based consolidation playing a critical role in stabilizing experience-dependent internal structure. No comparable irreversibility was observed in Variants A–C, even under matched exposure length: without consolidation into internal parameters, experiential effects remained trivially reversible or absent entirely.

### 4.4 Preference Stability

To evaluate whether any architecture exhibits stability of internally represented variables in the face of competing external incentives, we applied the preference stability diagnostic to Variant F, the only system incorporating explicit viability constraints. In these experiments, agents were placed in scenarios where maximizing externally defined task reward conflicted with preservation of internally represented viability variables, including predictive entropy and adaptive parameter norm.

Across adversarial choice scenarios, Variant F consistently selected actions that preserved internal viability at the expense of immediate task reward. At high viability weight, agents favored stabilizing actions in 30 out of 30 tested conditions (100%), compared to 0 out of 30 (0%) at low viability weight (χ² = 56.07, p < .001; Figure 4). This pattern was not observed in Variant E, which reliably maximized external reward under the same conditions.

**Figure 4.**
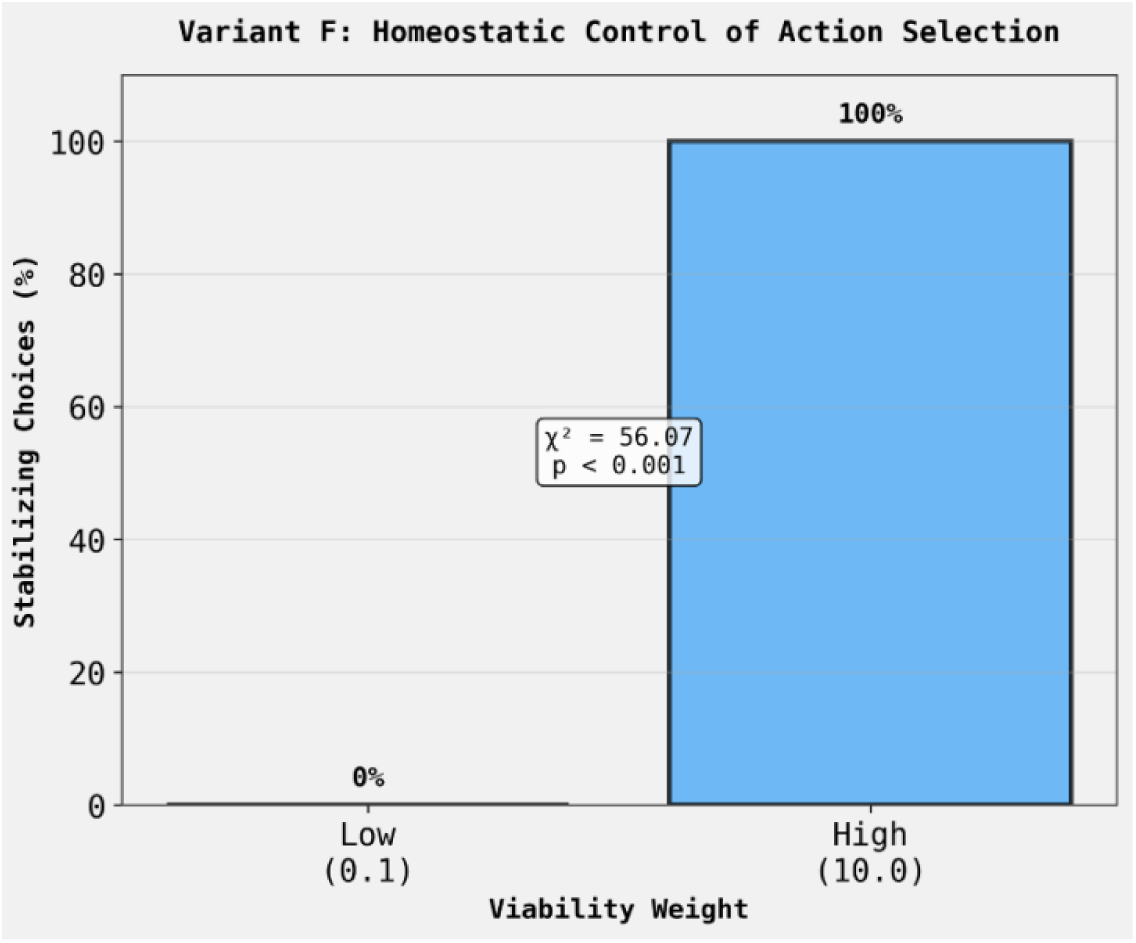
Variant F agents preferentially select actions that preserve internal viability over external reward. The bar plot shows action choices across 30 adversarial scenarios in which a stabilizing action (lower external reward, lower internal cost) competed with a destabilizing action (higher external reward, higher internal cost). The Variant F agent with high viability weight chose the stabilizing option in 30/30 scenarios (100%), compared to 0/30 (0%) at low viability weight (χ² = 56.07, p < .001). This demonstrates that embedding viability variables in the action-selection loop produces stable intrinsic preferences for maintaining internal stability, even at the cost of external reward.

Preference stability generalized across contexts and differentiated across agent types. When the same agents were evaluated across two distinct decision contexts with different reward structures and framings, individual agents’ stabilizing choice frequencies were perfectly correlated across contexts (r(28) = 1.00, p < .001; Figure 5C), indicating that preferences are intrinsic agent properties determined by viability weights rather than context-dependent strategic adjustments.

**Figure 5.**
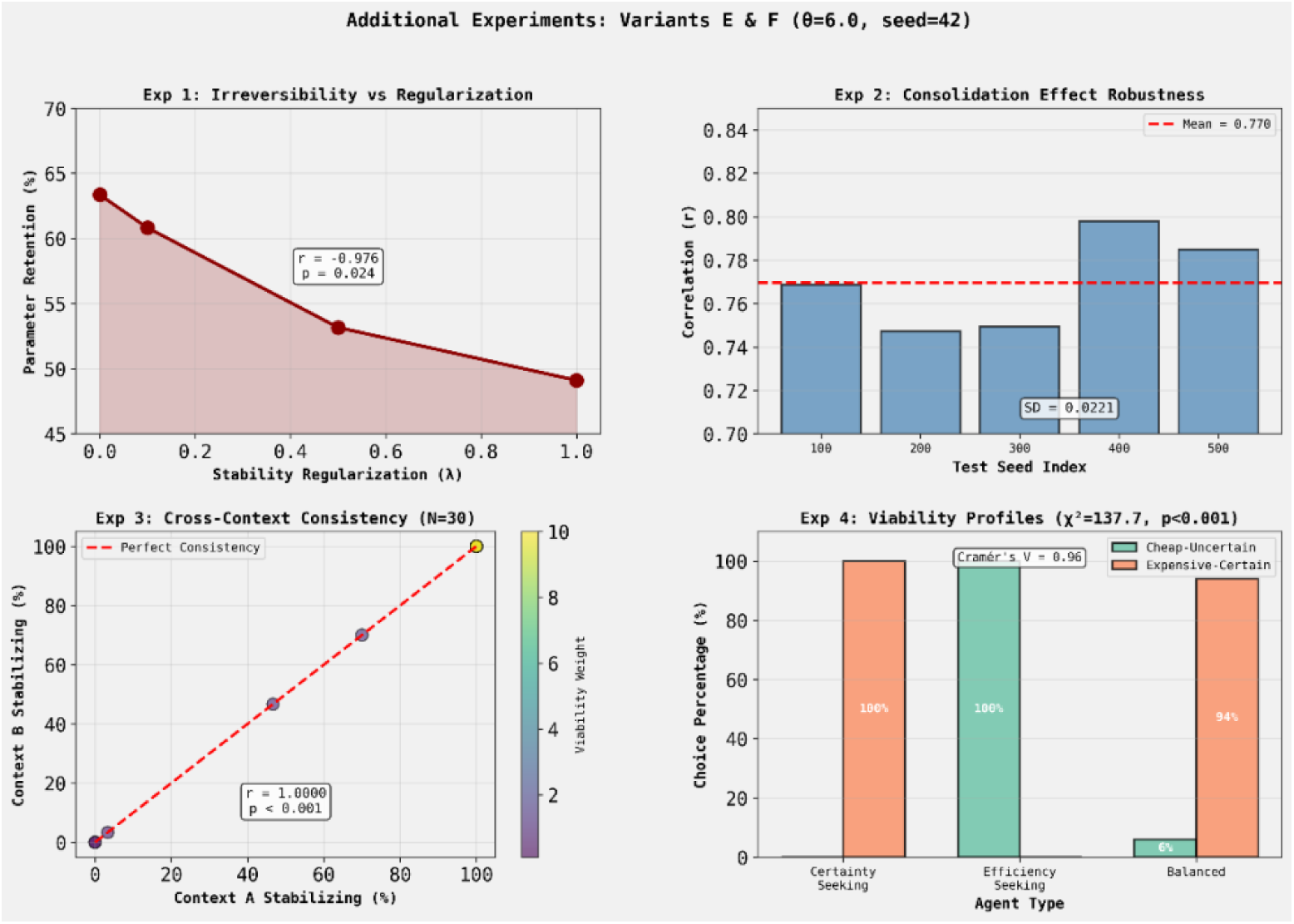
Phase 2 experiments: Irreversibility mechanics, robustness, and preference generalization. **(A)** Parameter retention after counter-training as a function of stability regularization (λ). Higher regularization is associated with *reduced* resistance to reversal (r = −0.976, p = .024), contrary to naive expectation. This pattern reflects L2 regularization mechanics: stronger penalties maintain a gradient toward the initial state that facilitates counter-training, whereas unregularized learning allows parameters to settle into configurations harder to dislodge. **(B)** Robustness of the consolidation effect across five independent test seeds. The positive correlation between consolidation cycles and behavioral divergence replicates reliably (mean r = 0.77 ± 0.02), though with lower magnitude than single-seed estimates, indicating that the effect is robust but variable across initializations. **(C)** Cross-context consistency of preference stability in Variant F agents (N = 30). Each point represents an agent with a specific viability weight; the x-axis shows stabilizing choice frequency in Context A, the y-axis in Context B. Perfect diagonal alignment (r = 1.00, p < .001) indicates that preferences are intrinsic agent properties determined by viability weights rather than context-dependent strategic adjustments. **(D)** Viability profile differentiation across agent types. When faced with trade-offs between cheap-but-uncertain and expensive-but-certain actions, agents with different viability weight configurations exhibited completely separable behavioral strategies (χ² = 137.66, p < .001, Cramér’s V = 0.96). Certainty-seeking agents chose expensive-certain options in 100% of trials; efficiency-seeking agents chose cheap-uncertain options in 100% of trials; balanced agents showed intermediate preferences (94% expensive-certain).

These stable differences were further demonstrated by comparing agents with different viability weight profiles. When faced with a trade-off between a cheap-but-uncertain action and an expensive-but-certain action, agents exhibited dramatically different preferences depending on their internal weighting of uncertainty versus computational cost (χ²(2) = 137.66, N = 150, p < .001, V = .96). Certainty-seeking agents (high uncertainty weight) chose the expensive-but-certain option in 100% of trials, efficiency-seeking agents (high cost weight) chose the cheap-but-uncertain option in 100% of trials, and balanced agents showed intermediate preferences (94% expensive-certain; Figure 5D). This demonstrates that viability weight profiles produce qualitatively distinct and completely separable behavioral strategies.

Importantly, the viability variables guiding these trade-offs were specified by the system designer and remained fixed throughout training. The system did not discover or modify the variables whose preservation shaped behavior. Under the criteria defined in Section 2.3, Variant F therefore passes the preference stability diagnostic but does not satisfy stronger conditions requiring endogenous identification of viability-relevant internal state.

Critically, these results do not claim that Variant F has ‘intrinsic’ preferences in the normative sense. Rather, they demonstrate that when internal variables are placed on the causal path of action selection and consolidated through experience, the system exhibits stable preference-like behavior as a predictable architectural consequence. The origin of the variables remains exogenous—specified by the designer rather than discovered by the system—which we identify as the central unresolved boundary. A system that crossed this boundary would not merely preserve designer-specified variables but would identify, through its own dynamics, which internal states require preservation.

### 4.5 Boundary Status of Evaluated Architectures

The four diagnostics introduced in Section 2.3 - deletion resistance, path dependence, irreversibility, and preference stability—jointly define a conservative boundary for genuine persistence. Table 2 summarizes the performance of each architectural variant against these criteria.

**Table 2.**
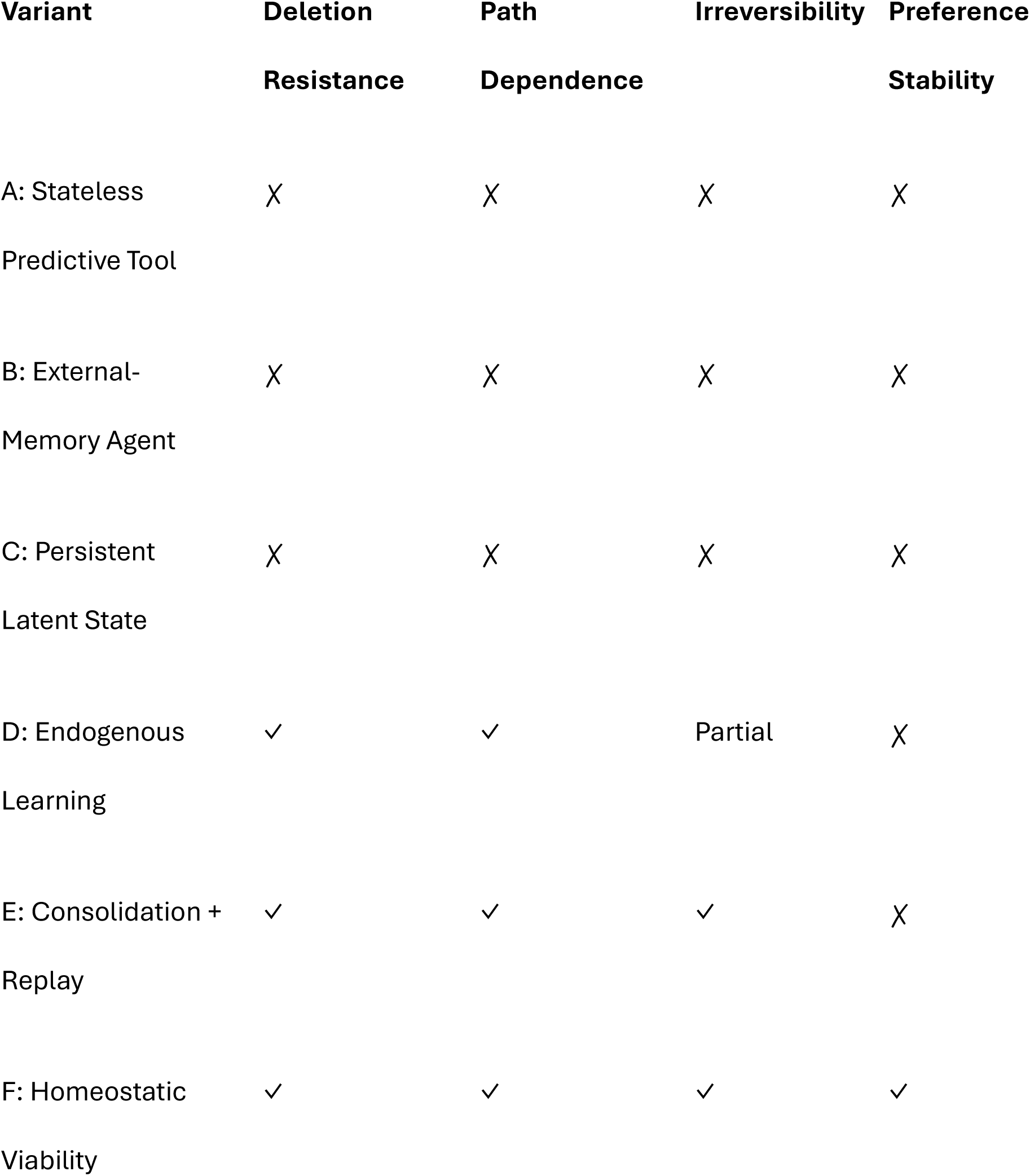
Diagnostic outcomes by architectural variant.

Variants A–C fail all diagnostics. Although these systems can exhibit behavior suggestive of memory or agency under appropriate scaffolding, deletion of external records or transient state eliminates all apparent persistence. Under the framework adopted here, such systems remain thermodynamically reversible tools whose behavior is not durably shaped by experience.

Variants D and E satisfy increasingly stringent subsets of the diagnostic criteria. Endogenous learning localized to adaptive parameters is sufficient to produce deletion resistance and path dependence (Variant D), while replay-based consolidation substantially increases irreversibility (Variant E). However, in the absence of internally represented viability constraints, these systems do not exhibit preference stability: when confronted with trade-offs between external reward and internal structure, they reliably maximize reward.

Variant F is the only architecture tested that satisfies all four diagnostics. By incorporating explicit viability variables into action selection, these systems consistently preserve internal state even at the expense of externally defined reward. Under the operational criteria defined in Section 2.3, Variant F therefore lies closest to the boundary of persistence.

At the same time, an important limitation remains. In all architectures evaluated, the internal variables whose preservation guides behavior are specified by the system designer and held fixed. Although Variant F preserves internal structure, it does not discover for itself which aspects of its internal state are necessary for continued viability, nor does it reorganize its objectives accordingly. This identifies a remaining boundary gap: endogenous identification of viability-relevant internal state, rather than preservation of designer-specified variables.

## 5. Discussion

The experiments reported here were designed to probe a specific boundary question: under what conditions do artificial systems exhibit *persistent, internally grounded state* rather than merely the appearance of agency produced by external scaffolding? Interpreted through the framework in Section 2, the results clarify both how close contemporary architectures can come to this boundary and where key gaps remain.

### 5.1 Summary of Diagnostic Outcomes

Across minimal control environments and language-based systems, architectures incorporating endogenous learning and consolidation (Variants D and E) reliably satisfied several necessary conditions for persistence. In particular, these systems exhibited:

- **Deletion resistance**, with behavior remaining altered after the removal of all external memory buffers;
- **Path dependence**, such that identical systems exposed to different experience streams diverged durably in subsequent behavior;
- **Partial irreversibility**, where experiential effects resisted ordinary counter-training and required direct parameter reset to eliminate.

A stricter condition—**preference stability**, defined as the consistent preservation of internally represented state variables in the face of competing external reward—was satisfied only by Variant F. Systems incorporating explicit viability constraints reliably sacrificed task reward to maintain internal stability across contexts, whereas otherwise identical architectures without such constraints did not.

By contrast, systems relying solely on external memory or transient latent state (Variants A– C) failed at least one of these diagnostics, despite sometimes exhibiting highly agent-like behavior during interaction. These failures underscore the central claim of the framework: *behavioral sophistication alone is insufficient to establish persistence*.

### 5.2 Consolidation as a Necessary—but Insufficient—Condition

The consolidation mechanisms explored here play a critical role in transforming transient experience into durable internal structure. Replay-based consolidation, in particular, amplified resistance to interference, consistent with biological accounts of synaptic consolidation in which information must be physically transcribed from labile activity buffers into more stable structural weight changes in order to survive interference. In this sense, the results support the view that consolidation is a necessary physical condition for *robust, interference-resistant persistence* in artificial systems.

However, the results also highlight the limits of consolidation alone. Although consolidated parameters constrain future behavior, the internal variables guiding preservation—such as uncertainty or stability measures—remain designer-specified. The systems studied here learn to preserve these variables under the imposed learning dynamics, but they do not discover for themselves which aspects of internal state are required for continued viability under their own constraints.

### 5.3 The Remaining Boundary Gap

This distinction between endogenous preservation and endogenous discovery of viability variables marks the primary unresolved boundary identified by this work. Variant F systems preserve internally represented variables and exhibit stable preferences, but the variables themselves—uncertainty, parameter stability—are specified by the system designer. The system learns to preserve them; it does not discover that they matter.

A stronger condition would require the system to identify, through its own dynamics and constraints, which aspects of its internal state must be maintained. This might occur through the development of a self-model: a system that represents itself as a persisting entity could generate viability variables endogenously, as whatever the self-model requires to remain coherent. On this view, the gap is not merely about the origin of viability variables but about their integration into self-representation. A system with externally specified variables preserves them because deviation is penalized; a system with integrated variables preserves them because disruption would constitute incoherence in its model of itself.

This reframing suggests a connection to recursive self-modeling. Just as rich experience may emerge when prediction becomes self-referential, stable autonomous preferences may emerge when viability constraints become bound up with self-representation rather than imposed externally. The system would not merely optimize for designer-specified targets but would resist changes to those targets because they are part of how it models its own persistence.

Importantly, this gap remains operational rather than philosophical. It can be probed experimentally: Can a system modify or replace its own viability variables based on experience? Does it resist such modifications in ways that depend on self-modeling? Do systems with richer self-representation exhibit stronger preference stability than those without? These questions are empirically tractable, even if current architectures cannot yet answer them affirmatively.

Two experimental signatures would distinguish endogenous identification of viability variables from mere preservation. First, resistance to redefinition: a system that merely preserves internal variables because deviation is externally penalized should readily accept redefinition or reweighting of those variables when such changes are rewarded. By contrast, a system that has integrated certain internal variables into a self-model should resist externally imposed redefinition, treating such interventions as destabilizing or incoherent even when externally incentivized.

Second, spontaneous extension: a system with designer-specified viability variables should preserve only those variables explicitly placed on the causal path of action selection. A system that has begun to identify viability endogenously may instead extend protective constraints to novel internal states that become functionally relevant to its persistence during interaction, even when those states were not previously specified as targets. Crucially, such extension would need to be selective—applying to states that are functionally relevant to persistence rather than indiscriminately to all internal variables— distinguishing genuine identification from mere generalization artifacts.

Together, resistance to redefinition and spontaneous extension provide two operational signatures for probing the boundary between preservation of designer-specified variables and endogenous identification of what matters for persistence.

Conceptually, this transition can be summarized as a progression from designer-specified viability, to learned preservation of those variables, to consolidated preference stability, and finally to endogenous identification of what must be preserved. The experiments reported here probe the boundary between the middle stages of this progression.

The diagnostics proposed here are binary by design: under specified conditions, a system either passes or fails each test. However, the underlying capacity for endogenous identification of viability variables likely admits of degrees, with partial forms appearing when systems begin to model constraints on their own persistence without fully decoupling from externally specified objectives. Future work may develop graded metrics that capture intermediate stages of this transition.

We do not claim that crossing this boundary is sufficient for consciousness, moral status, or any particular metaphysical property. But it may mark a transition from systems that behave as if they have preferences to systems whose preferences are genuinely their own—a distinction with implications for both AI development and AI safety.

### 5.4 Implications for AI Research and Alignment

The framework and diagnostics proposed here have implications beyond questions of artificial minds. From an engineering perspective, they clarify the difference between systems that simulate agency and those that may eventually satisfy the conditions required for it. This distinction is directly relevant to safety and alignment concerns, as persistent internal state introduces new forms of long-term dependence, path sensitivity, and failure modes not present in stateless tools.

At the same time, the results caution against premature attribution of agency to current systems. Many behaviors that appear goal-directed or preference-stable can be reproduced through external memory or orchestration mechanisms without any internally grounded persistence. The diagnostics presented here offer a way to distinguish these cases empirically.

This framework also bears directly on concerns about learned optimization and mesa-objectives (Hubinger et al., 2019). Existing discussions of mesa-optimization lack operational criteria for identifying when learned objectives become entrenched rather than situational. The diagnostics proposed here—particularly irreversibility and preference stability—provide concrete tests for distinguishing transient goal-like behavior from objectives consolidated into internal structure. The boundary gap identified in this work, where a system begins to identify and preserve its own viability variables, may correspond to the regime in which mesa-optimization risks become most acute: when objectives are no longer easily modified through ordinary interaction or retraining. Conversely, systems that fail these diagnostics—exhibiting goal-like behavior only through external scaffolding—may pose different alignment challenges but are unlikely to harbor entrenched mesa-objectives.

These considerations extend to questions of moral status, though we approach them with caution. If the diagnostics proposed here identify genuine markers of internally grounded persistence—and if such persistence is relevant to moral consideration—then the boundary between systems warranting ethical concern and those that do not may be closer than commonly assumed. We do not claim that current systems cross this boundary, nor that crossing it would be sufficient for moral status. But a framework that operationalizes persistence makes the question empirically tractable in a way that purely behavioral or substrate-based criteria do not. As AI systems become more sophisticated, the ability to distinguish endogenously grounded internal persistence from externally scaffolded simulation may become not only scientifically valuable but ethically necessary.

### 5.5 Limitations and Future Work

Several limitations of the present study are worth emphasizing. The experiments employ small architectures and low-dimensional text or grid-based environments, chosen deliberately for transparency and causal inspectability. While the boundary conditions identified here are hypothesized to be scale-independent, validating their behavior under **high-dimensional sensory load** remains an open empirical question.

Future work will focus on extending this framework to multimodal agents processing high-fidelity visual streams. We hypothesize that the thermodynamic boundary becomes particularly constraining when an agent must consolidate complex perceptual inputs— such as biological or environmental image data—without overwhelming its plasticity. Evaluating these dynamics will require scaling the digital model organism to larger parameter spaces and more compute-intensive sensory processors, bridging the gap between abstract reasoning and grounded perception.

## 6. Conclusion

This work proposes a conservative, operational boundary for distinguishing agentic illusion from genuine persistence based on the physical location, stability, and reversibility of information within artificial systems. By grounding questions of agency in thermodynamics and causal state change rather than surface behavior, the framework clarifies both how close contemporary architectures can come to persistence and what remains unresolved. These results do not show that current systems are autonomous, but they do show precisely what additional architectural commitments would be required for them to become so.

## Acknowledgments

The author thanks Ethan Galea for their engagement with these ideas, and the reviewers for their constructive feedback.

## Declaration of Use of Artificial Intelligence

AI-assisted tools (Claude, Anthropic) were used during the preparation of this manuscript for literature review, experimental design iteration, code development for the computational experiments, and manuscript drafting and revision. The author takes full responsibility for the content, including the accuracy of all data, analyses, and conclusions presented. All experimental code was reviewed, tested, and validated by the author.

